# Identification of Raptor and GLI1 as USP37 substrates highlight its context-specific function in medulloblastoma cells

**DOI:** 10.1101/2025.04.22.650057

**Authors:** Ashutosh Singh, Donghang Cheng, Amanda Haltom, Yanwen Yang, Tara Dobson, Rashieda Hatcher, Veena Rajaram, Vidya Gopalakrishnan

## Abstract

*USP37* gene encodes a deubiquitylase (DUB), which catalyzes the proteolytic removal of ubiquitin moieties from proteins to modulate their stability, cellular localization or activity. Its expression is downregulated in a subgroup of medulloblastomas driven by constitutive activation of sonic hedgehog (SHH) signaling. Patients with SHH-driven medulloblastomas with elevated expression of the *RE1* silencing transcription factor (REST) and reduced expression of *USP37* have poor outcomes. In previous studies, we showed sustained proliferation of SHH-medulloblastoma cells due to blockade of terminal cell cycle exit and neuronal differentiation stemming from a failure in USP37-dependent stabilization of its target, the cyclin-dependent kinase inhibitor (CDKI)-p27. This finding suggested a tumor suppressive function for USP37. Interestingly, the current study also uncovered Raptor, a component of the mTORC1 complex, as a novel target of USP37. Under conditions of low-*USP37* expression, reduced Raptor stability and mTORC1 activity caused a decline in phosphorylation of 4E-binding protein 1 (4EBP1) and increased its interaction with eukaryotic elongation factor 4E (eIF4E), which is known to inhibit CAP-dependent translation initiation. Surprisingly, a subset of patients with SHH-driven medulloblastomas with elevated expression of *USP37* and the Glioma associated Oncogene 1 (*GLI1*), also exhibited poor outcomes. Using genetic and biochemical analyses, we showed that USP37-mediated stabilization of GLI1, a terminal effector of SHH signaling, increases pathway activity and upregulates expression of its target oncogene products, NMYC and CCND1, to drive cell proliferation. These data indicate that USP37 elevation in SHH-driven medulloblastomas has the potential to promote non-canonical activation of SHH signaling. Overall, our findings suggest that USP37 may have context-specific oncogenic and tumor suppressive roles in medulloblastoma cells.

## Introduction

Medulloblastoma (MB) is the most common malignant brain tumor of childhood, with over 500 new diagnoses each year [1–8]. Despite advances in the molecular stratification and our understanding of tumor biology, standard of care relies on surgical resection, craniospinal radiation and chemotherapy [7, 9–19]. These are toxic to the normal developing brain and cause long-term quality of life issues and further, are ineffective in the setting of recurrence/metastasis [9–15, 20]. The 5-year survival for patients diagnosed with MBs is 66%, with their molecular subgroup - Wingless (Wnt), Sonic Hedgehog (SHH), group 3, or group 4, serving as a main determinant of outcomes [1, 3, 8, 9, 11, 21–33]. Patients with group 3 MBs have the worst prognosis, whereas those with SHH and group 4 tumors have intermediate outcomes [1, 3, 8, 9, 11, 21–33]. While deregulated chromatin remodeling is implicated in MB genesis [21, 26, 32, 34–49], there is also growing evidence for aberrant regulation of protein degradation in SHH-driven MBs [35, 36].

In previous work, we implicated Ubiquitin Specific Protease 37 (USP37) - a deubiquitylase (DUB) and a component of the proteasomal degradation machinery, in MB biology [35, 36]. DUBs catalyze the proteolytic removal of ubiquitin groups from proteins, causing changes in their stability, sub-cellular localization, or function [50–54]. They oppose the functions of enzymes (E1-3), which coordinate the addition of ubiquitin moieties to proteins [55–57]. In SHH-driven MB cells with *RE1* silencing transcription factor (*REST)* elevation, we demonstrated that *USP37* downregulation prevented the stabilization of the cyclin-dependent kinase inhibitor (CDKI)-p27 and maintained cell proliferation [35]. Pharmacological targeting of G9a histone methyltransferases upregulated *USP37* expression and reduced tumor formation of low-*USP37* MBs, suggesting an epigenetic basis for *USP37* downregulation in MBs [36]. Just prior to our report, a different group identified a role for USP37 in controlling S phase entry through its regulation of cyclin A stability in non-neural cells [58]. Since these initial reports, work from other groups has also shown the importance of USP37 in the regulation of DNA replication and cell cycle [35, 59, 60] / cell proliferation [61]in addition to functions in receptor signal transduction [62], homologous recombination [63], DNA damage repair [64], cell migration [65], autophagy [66], epithelial to mesenchymal transition [67] and chemotherapy resistance [68]. [69]

In the current study we identify Raptor and GLI1 as novel USP37 targets in SHH-MBs and also suggest that *USP37* may have context specific oncogenic roles in addition to its previously described tumor suppressive function in these tumors [35]. Based on our recent finding that *USP37* loss in the context of high-*REST* expression, and *USP37* elevation in the background of high-glioma associated oncogene 1 (*GLI1)* levels were both found to be correlated with poor outcomes for patients with SHH-MBs. We show that low USP37 results in lowered stability of Raptor, a component of the mTORC1 complex, and a regulator of protein translation. In the context of high *USP37* expression, we found increased GLI1 protein levels and USP37-dependent GLI1 ubiquitination and protein stabilization. These data suggest that USP37 elevation in SHH-driven MBs has the potential to promote non-canonical activation of SHH signaling.

## Materials and Methods

### Patient samples

Two publicly available MB patient’s datasets (GSE85217 and GSE124814) were used for gene expression analysis [70, 71]. Differential gene expression analysis was performed using the R2: Genomics Analysis and Visualization Platform (http://r2.amc.nl). A value of p< 0.05 was considered as significant. Gene ontology p-values are not corrected for multiple testing. Immunohistochemistry for REST and USP37 was performed in paraffin embedded de-identified MB tumor sections (n=33) by our collaborating neuropathologist.

### Cell culture

293T and SHH-MB cell lines (DAOY, UW228, UW426, and ONS76) are used in this study. DAOY cells were purchased from the American Type Culture Collection (ATCC) (Manassas, VA). UW228 and UW466 cells were a kind gift from Dr. John Silber at the University of Washington. ONS76 cells were purchased from Accegen, NJ, USA. All the cell lines were maintained in Dulbecco’s modified Eagle’s medium (Sigma-Aldrich, MO, USA), supplemented with 10% fetal bovine serum (Sigma-Aldrich), 1% antibiotic-antimycotic (Thermo Fisher Scientific, MA, USA), and 1% sodium pyruvate (Thermo Fisher Scientific) and grown at 37 °C with 5% CO2.

### Patient-derived xenograft models

Patient-derived xenograft (PDX) models (RCMB-18, 24 and 54) were a kind gift of Dr. Robert Wechsler-Reya (Columbia University). Serial transplantation of tumors was carried out in NOD.Cg-*Prkdc^scid^ Il2rg^tm1Wjl^*/SzJ (NSG) mice (Jackson Laboratory, Bar Harbor, ME), by intracranial inoculation of tumor cells using a stereotactic device as described previously [48]. Housing, maintenance and experiments involving mice were done in compliance with a protocol approved by the University of Texas MD Anderson Cancer Center’s Institutional Animal Care and Use Committee (IACUC). Our study is reported in accordance with ARRIVE guidelines (https://arriveguidelines.org).

### Animals

Generation of *Ptch^+/−^/REST^TG^* was described earlier [48]. The *hREST* transgene expression was induced by intraperitoneal (ip) injections of (100 μl of 2 mg/ml) tamoxifen (Cat# T5648, Sigma-Aldrich) on post-natal (P) days 2, 3 and 4. Moribund animals were euthanized, and brains were harvested for further analysis.

### Immunohistochemistry

Mouse brain tissues were fixed in 10% buffered formalin phosphate for 48 hours (h) and embedded in paraffin. 8µm thick sections were used for histological analysis using a Gemini AS Automated Stainer (Thermo Fisher Scientific) After overnight incubation with primary antibody at 4°C, the sections were incubated with biotinylated secondary antibody provided by either the ABC kit or the MOM kit (Vector Laboratories, CA, USA). For detection, VECTASTAIN® Elite® ABC-HRP Kit, Peroxidase (Cat# PK-6101, Vector Laboratories) were used according to the manufacturer’s instructions and developed using the DAB Substrate Kit, Peroxidase (HRP) (Cat# SK-4100, Vector Laboratories) followed by counterstaining with hematoxylin. After dehydration and mounting, slides were visualized under a Nikon ECLIPSE E200 microscope mounted with an Olympus SC100 camera. The list of primary antibodies used for IHC is provided in Supplementary Table S1.

### Lentiviral infection and generation of stable cell lines

Embryonic kidney (HEK) 293T cells were co-transfected with either a control or the gene of interest along with packaging plasmid (PAX2) and envelope plasmid (MD2). Lentiviral particles were collected 48-hours post-transfection. MB cells were transduced with the collected viral supernatant in the presence of Polybrene (8 μg/mL) and incubated for 48 hours. Infected cells were then cultured in medium containing 2 μg/mL puromycin for up to 1 week for selection.

### Western blot analyses

Cell lysates were prepared in lysis buffer (50 mM Tris-HCl pH 8.0, 50 mM NaCl, 1% NP-40, 0.5% sodium deoxycholate, 0.1% SDS, and protease/phosphatase inhibitors) and processed for Western blotting as described previously [40] using primary antibodies listed in Supplementary Table S1 followed by HRP-conjugated goat anti-mouse or anti-rabbit secondary antibodies. SuperSignal West Dura Extended Duration Substrate (Cat#34075, Thermo Fisher Scientific) and Western Lightning Plus-ECL, Enhanced Chemiluminescence Substrate (Cat# 50-904-9325, Fisher Scientific, MA, USA) were used to develop the blot and detected using Kodak Medical X-Ray Processor 104 (Eastman Kodak Company) or ChemiDoc Touch Imaging System (Bio-Rad). Images were analyzed using Image Lab software version 5.2.1 (Bio-Rad).

### Co-immunoprecipitation

Cell pellets were washed with ice-cold PBS and lysed in mild lysis buffer (50mM Tris–HCl pH 7.5, 150mM NaCl, 1mM EDTA, 1% Triton X-100, 5 mM EDTA) containing protease inhibitor cocktail (Cat# 78429, Thermo Fisher Scientific) and phosphatase inhibitor (Sigma) and sonicated. Lysates were incubated with control mouse IgG, anti-USP37 (Cat# A300-927A, Thermo Fisher Scientific), anti-4EBP1 (Cat# 9644, Cell Signaling Technology, MA, USA), anti-eIF4E (Cat# 2067, Cell Signaling Technology) primary antibodies overnight at 4°C and then incubated with Pierce™ Protein A/G UltraLink™ Resin (Cat# 53132, Thermo Fisher Scientific) for 1.5 hr at 4°C. After four washes with lysis buffer, beads were boiled in loading buffer, separated by SDS-PAGE, transferred onto PVDF membranes, and analyzed by Western blotting.

### In vitro deubiquitination (DUB) assay

293T cells were co-transfected with pcDNA3-myc3-Raptor/ pcDNA3-6XHis-GLI1 and HA-Ub or with pDEST26-FLAG-HA-USP37, FLAG-HA-USP37^C350S^ or FLAG-HA-USP1 using jetPRIME (Polyplus, NY, USA). Cells were treated with 20mM MG132 for 6h prior to lysis. HA-Ub-Myc-Raptor and HA-Ub-6XHis-GLI1 substrate were purified using EZ view Red Anti-c-Myc Affinity Gel (Cat# E6654, Millipore Sigma) and Ni-NTA Agarose (Cat # R90101, Thermo Fisher Scientific), respectively. Deubiquitinase (DUBs: USP37, USP37^C350S^ or USP1) were immunopurified using anti-FLAG M2 beads (Cat# A2220, Millipore Sigma) and eluted with FLAG peptide (Cat# HY-P0319, MedChem Express, NJ, USA). For DUB assays, equal amounts of substrates and purified DUBs were incubated at 37°C in the presence or absence of 15mM N-ethyl maleimide (NEM). Reactions were terminated by boiling in 2X Laemmli buffer. Reactants were analyzed by Western blotting.

### In vivo DUB assay

MB cells were transiently transfected with plasmids expressing (*Flag-HA-USP37^WT^* and *Flag-HA-USP37^C350S^* (Addgene, MA, USA). Cells were treated with 20mM MG132 for 6h prior to lysis. Samples were heat denatured and subjected to SDS-PAGE and Western blotting using anti-Raptor (Cat# 2280, Cell Signaling Technology), anti-GLI1 (Cat# ab134906, Abcam, MA, USA), anti-FLAG (Cat# F1804, Millipore Sigma) and anti-Ubiquitin (Cat# 3933, Cell Signaling Technology) antibodies.

## Results

### Low- and high-USP37 expression in SHH-MBs is correlated with poor patient outcomes

Our previous work reported *USP37* as a novel target of REST in SHH-MBs and also studied its expression in MB cell lines compared to normal cerebella [35, 36]. Here, we assessed the levels of USP37 in human MB samples (n=33) (Fig. 1A). Samples were graded as negative (-) or positive (+/ ++/ +++/ ++++) for USP37 expression (Fig. 1A). Approximately, 33.3% of the samples were negative for USP37, whereas weak and focal staining (+) was noted in 30% of the samples. Weak diffuse and multifocal focal staining (++) was noted in 6.7% samples (Figs. 1A and 1B). The remaining 30% of samples exhibited strong expression of USP37 (23.3% - strong and focal staining (+++) and (6.7% - strong diffuse or focal (++++)) (Figs. 1A and 1B). Further, samples with negative or weak expression of USP37 (-/+/++) exhibited high levels of REST (++/+++/++++), while samples with high USP37 (+++/++++) were negative or expressed low levels of REST (-/+) (Fig 1C).

**Figure 1.**
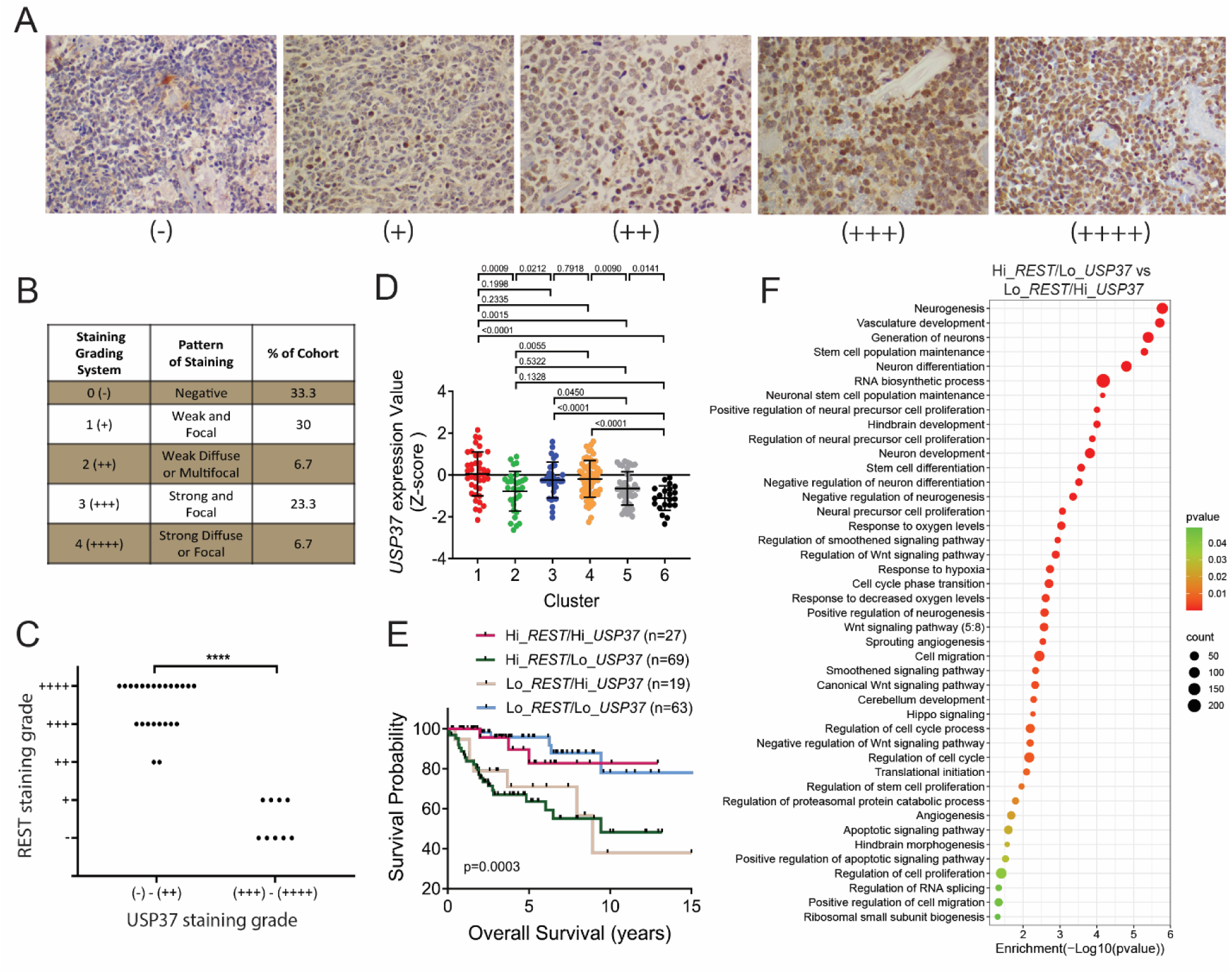
High- and low-*USP37* in SHH-driven MBs is associated with poor patient outcomes. **(A)** Human MB samples (n=33) were stained with anti-USP37 antibodies and graded on a scale from (-) to (++++) based on the level of USP37 expression (magnification: 40X). **(B)** Quantitation of data shown in (A) to show the distribution of tumors with different grades and patterns of USP37 staining. **(C)** Graphical representation of data from (A) and (B) to show the inverse correlation between REST and USP37 levels. Each dot represents a tumor. Significance and correlation were measured using the unpaired t-test and Pearson correlation coefficient (r= 0.478 and P value= <0.0001). **(D)** *USP37* gene expression in a microarray data set (GSE85217) [70] of human SHH- MB samples. Hierarchical clustering based on expression of neuronal differentiation markers divided the SHH-type MB patient samples into six distinct clusters [48]. Each dot corresponds to an individual patient. **(E)** Kaplan–Meier plot to demonstrate significant differences in the overall survival of SHH-MB patients divided into 4 cohorts based on the relative expression of *REST* and *USP37* in their tumors (GSE85217) [70]. **(F)** Pathway analysis to show significantly enriched pathways in tumors with high-*REST*/low-*USP37* expression relative to samples with low-*REST*/high-*USP37* expression using the GSE124814 dataset [71].

Transcriptomic data from MB patient samples revealed that *USP37* levels were significantly lower in WNT, SHH and group 3 tumors compared to group 4 samples (Fig. S1A). Since we had shown a role for USP37 in SHH-MB pathology previously [35], we further evaluated *USP37* mRNA in the six differentiation-based clusters of SHH-MB patients described in our prior study [48, 70]. As expected, clusters 2 and 5, previously described as having significantly elevated REST levels and poor outcomes also showed significantly reduced *USP37* expression (Fig. 1D). Kaplan Meier analyses confirmed that SHH-MB patients with high-*REST*/low-*USP37* (n=69) did indeed exhibit poor survival compared to patients with high-*REST*/high-*USP37* (n=27) and low-*REST*/low-*USP37* (Fig. 1E). Unexpectedly, a cohort of SHH-MB patients with low-*REST*/high-*USP37* (n=19) in their tumors also exhibited poor outcomes (Fig. 1E). Further evaluation of the high- *REST*/low-*USP37* and low-*REST*/high-*USP37* cohorts of patients from the GSE124814 and GSE85217 datasets by pathway analyses revealed a significant enrichment in processes known to regulate cerebellar and hind brain development, including those regulating proliferation and maintenance of stem-cells and neuronal precursor cells, neurogenesis, and neuronal differentiation, Smoothened, Wnt and Hippo signaling (Fig 1F and S1B). Interestingly, pathways which we previously discovered as REST-regulated such as vasculogenesis, angiogenesis, hypoxia, cell-cycle and cell migration were enriched in the high-*REST*/low-*USP37* cohort of patient samples compared to the low- *REST*/high-*USP37* group (Fig 1F and S1B). Apoptosis, protein translation, and mTOR were also differentially enriched in those two patient cohorts (Fig 1F).

### Animal models of SHH-MBs express variable levels of USP37 and REST

Immunohistochemical staining of patient-derived orthotropic xenograft (PDOXs) sections of SHH-MB tumors were performed to assess the levels of USP37 and REST (Fig 2A). Of the four PDOX tumors, two showed strong Ki67 staining, while all four showed some positivity for TUBB3. Strong REST staining was observed in all the four samples. Varying levels of USP37 staining were observed in the tumors, with RCMB-018 showing minimal staining and Med1712 exhibiting maximum cytoplasmic USP37 expression among the four samples (Fig 2A). Cerebellar sections of *Ptch^+/-^/REST^TG^* animals bearing tumors were also histologically analyzed, which revealed strong REST staining as expected (Fig 2B). USP37 was low in two of the three samples studied (Fig 2B).

**Figure 2.**
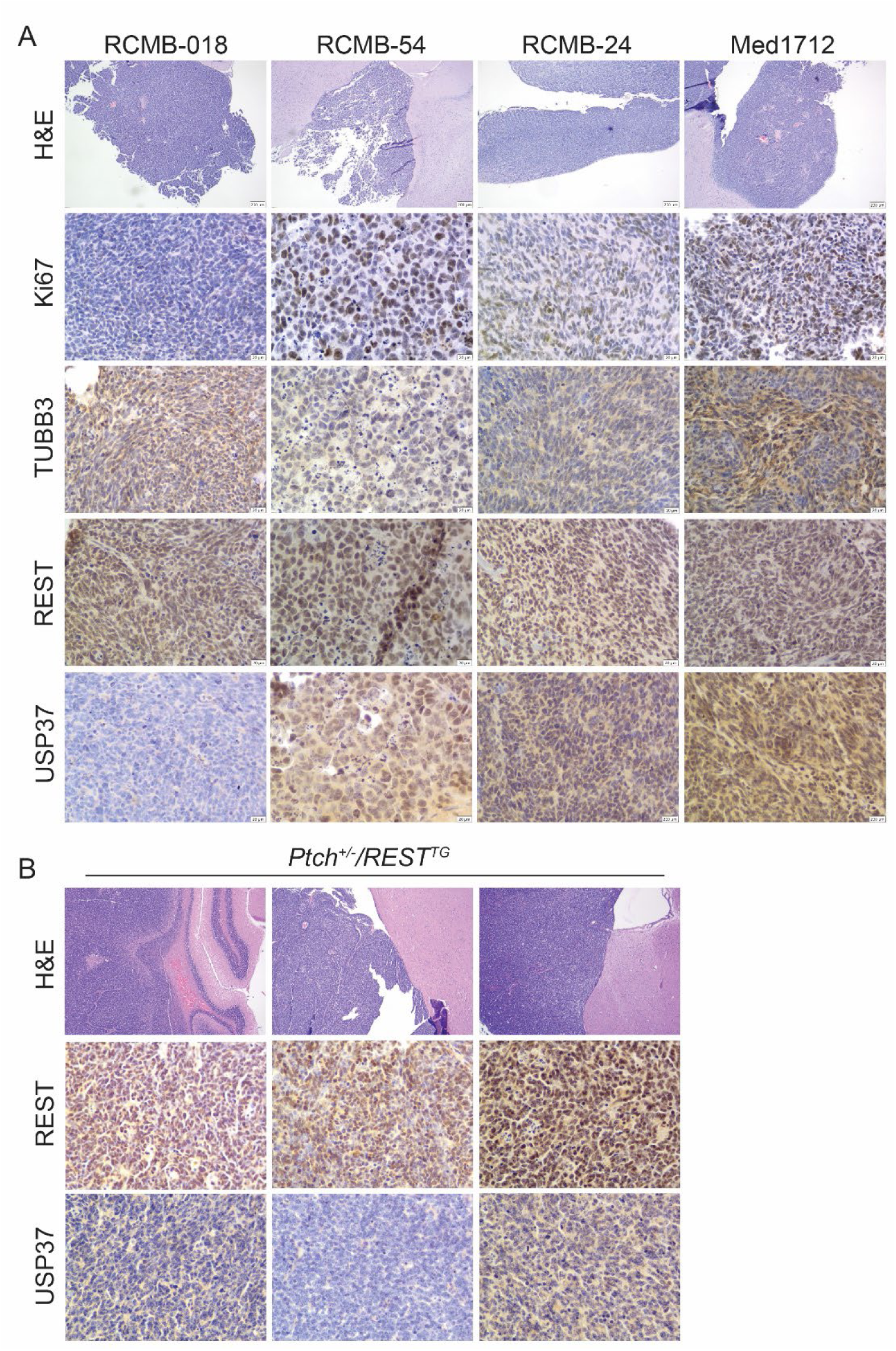
H&E and IHC analyses of mouse models of SHH-driven MBs. Staining of cerebellar sections from **(A)** mice with patient-derived orthotopic xenografts and **(B)** tumor-bearing *Ptch^+/−^/REST^TG^* animals (n=3) for Ki67, TUBB3, REST, USP37, and REST and Usp37, respectively (scale bar: 200µm for H&E and 20µm for IHC).

### Raptor is a novel target of USP37

In previous work by Dobson et al, we found that REST elevation in SHH-MB tumors to correlate with AKT activation as measured by its phosphorylation at Ser473 (pAKT^Ser473^), a modification brought about by the mammalian target of rapamycin complex 2 (mTORC2) [48]. Western blotting (Fig. 3A) confirmed these findings in human SHH-MB cell lines. Cells with higher levels of REST (293 and DAOY) had higher p-AKT^Ser473^ levels and that of its known targets p-GSK-3b^Ser9^ and p-p27^Thr157^ compared to lower REST expressing cells (UW228 and UW426) [72, 73] (Fig. 3A). Total GSK-3b and p27 were included as controls (Fig. 3A). p27 levels were lower in DAOY cells consistent with our previous report demonstrating that reduced USP37 levels prevented the stabilization of the protein in these cells [35]. It also suggested that the kinase activity of Rictor was higher in 293 and DAOY cells compared to UW228 and UW426 cells as measured by reduction in the levels of the Thr1135 phosphorylation, an event known to inhibit Rictor activity [74, 75] (Fig. 3B). Total Rictor as well as MLST8 levels were not significantly different in these cells (Fig. 3B). Interestingly, mTOR expression was lower in UW228 cells whereas expression of mSIN1 expression, a structural component of the mTORC2 complex, was lower in UW228 and UW426 cells compared to DAOY and 293 cells (Fig. 3B). b-actin served as the loading control (Fig. 3B). These results show that REST elevation is associated with increased mTORC2 complex activity in SHH-MB cells.

**Figure 3.**
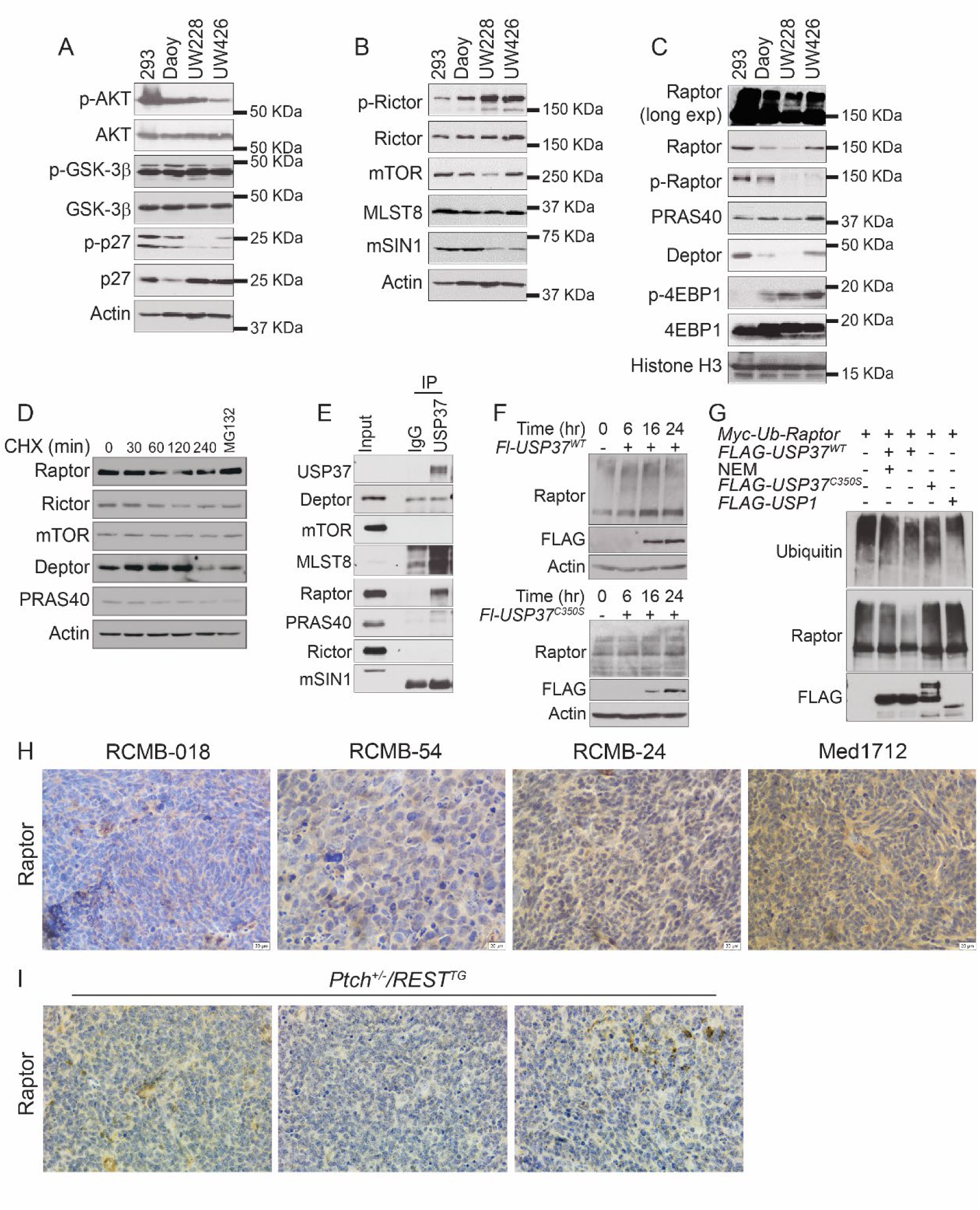
Raptor is a target of USP37. **(A)** Western blot to assess levels of AKT, GSK3β, and p27 and their respective phosphorylated forms in different cell lines with varying levels of REST. Western blot to study levels of **(B)** mTORC2 components -p-Rictor, Rictor, mTOR, mLST8, and mSIN1 in SHH-MB cell lines with varying levels of REST, and **(C)** mTORC1 components - Raptor p-Raptor, PRAS40, Deptor, and the mTORC1 target - 4EBP1 and its phosphorylated form. **(D)** Western blotting to demonstrate post- translation and proteasomal control of mTORC1/2 components (Raptor, Rictor, mTOR, Deptor, and PRAS40) following treatment with Cycloheximide (CHX) and without (lanes 1-5) or with MG132 (lane 6) in DAOY cells **(E)** Co-immunoprecipitation assay using control IgG or anti-USP37 antibodies followed by Western blotting to assess interaction between USP37 and mTORC1/2 components. Input lane shows 2% of total lysate. **(F)** *In vivo* DUB assay using transiently transfected FLAG (Fl)-tagged wildtype (WT) USP37 and mutant USP37 (USP37^C350S^) to study longitudinal changes in endogenous Raptor levels in DAOY cells **(G)** *In vitro* DUB assay was done by co-incubation of purified MYC tagged- Ub-Raptor with Fl-WT-USP37 in the presence of NEM (lane 2), Fl-WT-USP37 in the absence of NEM (lane3), Fl-USP37^C350S^ (lane 4) and Fl-USP1 (lane 5). The reaction containing MYC tagged-Ub-Raptor substrate alone is included as a control (lane 1). IHC analysis showing level of Raptor in cerebellar sections of mice with **(H)** human SHH-MB PDOX tumors and **(I)** *Ptch^+/−^/REST^TG^* tumors (scale bar: 20µm).

AKT signaling is also known to negatively control mTORC1 activity. Indeed, Western blot analyses showed lower mTORC1 activity in DAOY cells compared to UW228 and UW426 cells based on levels of phosphorylated 4EBP1, although total 4EBP1 levels were not different between these cells (Fig. 3C). Deptor, a component of both mTORC1 and mTORC2 complexes was absent in UW228 cells and lower in DAOY cells compared to 293 and UW426 cells (Fig. 3C). PRAS40 levels was similar between the cells (Fig. 3C). However, Raptor levels were significantly reduced in DAOY cells and were also associated with higher levels of the inhibitory Ser792 phosphorylation and a laddering migration pattern suggestive of post-translational modification such as ubiquitination [76, 77] The decrease in Raptor levels in DAOY cells following treatment with cycloheximide (CHX) suggested that its expression is post-transcriptionally controlled in SHH-MB cells (Fig. 3D). Further, treatment with the proteasome inhibitor, MG132, countered the reduction in Raptor levels suggesting that Raptor may be subjected to ubiquitin-mediated proteasome degradation (Fig. 3D). These findings were confirmed in ONS76 cells (Fig. S2A). Other mTORC1 components did not reveal a similar MG132-dependent change in levels (Fig. 3D). Immunoprecipitation assays were performed following co-transfection of DAOY cells with plasmids expressing *Myc-Raptor* and *HA-Ubiquitin*. Pull down with anti-HA antibodies HA revealed strong ubiquitination of Raptor (Fig S2B).

As a first step in investigating if the stability of Raptor is regulated by USP37, we performed co-immunoprecipitation assays using anti-USP37 antibody to demonstrate an interaction between endogenous USP37 and components of mTORC1 and mTORC2 in MB cells (Fig. 3E). Strong interaction between USP37 and Raptor was observed, but not between USP37 and other components of mTORC such as Rictor, Deptor, mTOR, pRAS40, and mSIN1 (Fig. 3E). ONS76 cells also exhibited a strong interaction between USP37 and Raptor (Fig. S2C). Next, we assessed the stability of Raptor in response to USP37 overexpression. *In vivo* DUB assays confirmed an increase in Raptor levels and a decrease in high-molecular weight bands of Raptor following USP37 overexpression in DAOY cells and ONS76 cells (Figs 3F and S2D). Raptor levels were not discernably different in cells overexpressing a catalytically inactive mutant of USP37 (USP37^C350S^) (Figs. 3F and S2D). We also performed *in vitro* DUB assay using purified tagged proteins to ask if USP37 directly deubiquitinates Raptor. Addition of USP37 to the reaction containing Myc-Ub-Raptor resulted in a substantial decrease in slower migrating forms of Raptor compared to that containing USP37 and a protease inhibitor N-ethyl maleimide or USP37^C350S^ or USP1 (Fig. 3G). Thus, our data thus far suggest that REST-dependent destabilization of Raptor involves *USP37* silencing. Immunohistochemical staining of SHH-MB PDOXs and *Ptch^+/-^/REST^TG^* tumor sections was also carried out to demonstrate a positive correlation between USP37 and Raptor levels (Figs. 2A, 3H and 3I). Together, these findings indicate Raptor is a novel target of USP37.

### REST and USP37 converge on the cellular cap-dependent protein translational machinery

*USP37* overexpression in MB cells-dependent increase in Raptor levels and targets downstream of it, was accompanied by an increase in phospho-4EBP1 (Fig. 4A). However, total 4EBP1 levels were unaffected (Fig. 4A). p27 levels were also increased under these conditions, as expected, and served as a positive control (Fig. 4A). Actin was used as a loading control (Fig. 4A). mTORC1 kinase controls protein translation initiation through phosphorylation of 4EBP1, which inhibits its binding with eIF4E and favors cap-dependent translation initiation (Fig. 4B) [78, 79]. However, as seen in Figs. 4A and 4C, *USP37* overexpression in DAOY and ONS76 cells caused an increase in phospho-4EBP1 levels, and co-immunoprecipitation assays using anti-4EBP1 antibodies showed an USP37-dependent decrease in interaction between 4EBP1 with eIF4E. An increase in the phosphorylated form of 4EBP1 was also noted in the immunoprecipitate, validating the decreased interaction between 4EBP1 and eIF4E and confirming increased activation of mTORC1 complex under conditions of USP37 elevation (Figs. 4A and 4C). As expected, 4EBP1 did not interact with eIF4G, a scaffolding component of the translational initiation machinery (Fig. 4C). These data suggest that cap-dependent translation initiation may be favored under conditions of constitutive *USP37* expression. Conversely, under conditions of REST overexpression and *USP37* downregulation, interaction between eIF4E and 4EBP1 was increased and that between IF4E and eIF4G was decreased, suggesting an inhibition of cap-dependent translation initiation (Fig. 4D).

**Figure 4.**
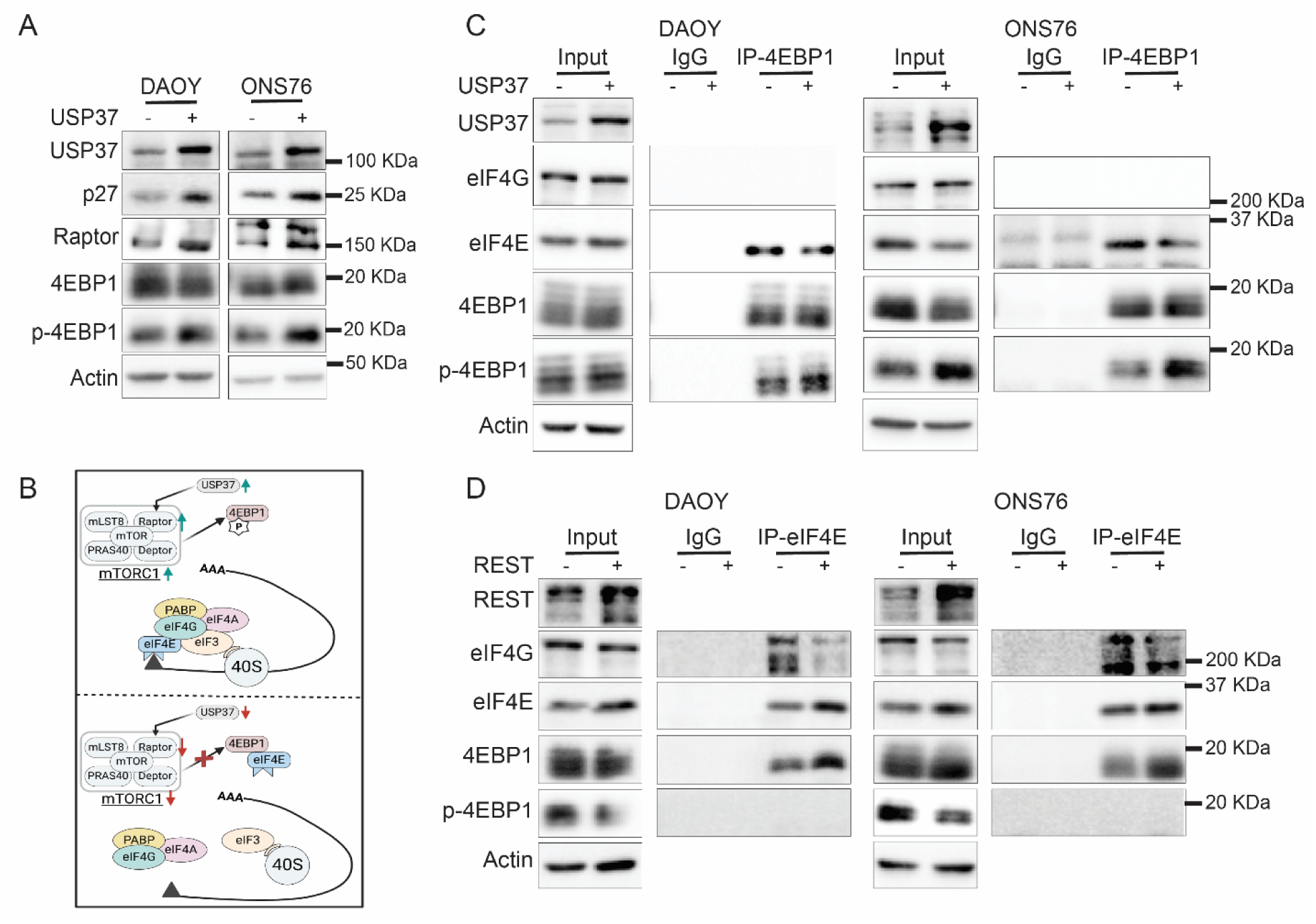
REST and USP37 modulate cap-dependent protein translation initiation machinery. **(A)** Western blotting to show changes in the levels of Raptor, 4EBP1, p- 4EBP1 and p27 with constitutive USP37 over expression in, DAOY and ONS76 MB cell lines. **(B)** Schema to illustrate expected effect of mTORC1 kinase activity on phosphorylation of 4EBP1 and its interaction with eIF4E. Phosphorylated 4EBP1 releases eIF4E to allow its binding to the CAP structure of *mRNA* allow canonical protein translation. Conversely, mTORC1 inactivity allows unphosphorylated 4EBP1 to bind to eIF4E and block its binding to the CAP structure of *mRNA* and prevent initiation of canonical protein translation, and potential shift to CAP-independent protein translation. Co-immunoprecipitation assays to study **(C)** interaction of 4EBP1 with eIF4G and eIF4E using control IgG or anti-4EBP1 antibody and **(D)** eIF4E with eIF4G, 4EBP1, and p- 4EBP1 using control IgG or anti-eIF4E antibody, in parental and isogenic USP37- and REST- overexpressing DAOY and ONS76 cells, respectively.

### USP37 promotes GLI1 stability

As shown in Fig. 1E, patients with high-*REST*/low-*USP37* as well as low-*REST*/high- *USP37* expression in their tumors exhibited poor survival. Enrichment of pathways associated with SHH signaling led us to investigate if the underlying mechanism may involve USP37-mediated regulation of one or more molecules driving SHH pathway activity. Based on a previous report that GLI1 is a target of USP37 in breast cancer cells - a non-neural cell type, we explored if a similar USP37-dependent stabilization of GLI1 and consequent hyperactivation of SHH signaling may drive SHH MB tumor growth [67]. Consistent with the above possibility, patients with high-*GLI1* expression, and low or high- *USP37* expression, in their tumors exhibited poor overall survival (Fig 5A). Pathways known to be controlled by GLI1 including apoptosis and cell migration, ciliary assembly, transport and organization, early-stage brain development, and development of metencephalon, hindbrain, cerebellum, and neuronal differentiation, were enriched in patient tumors with high GLI1 expression in the GSE124814 dataset (Fig 5B). Immunohistochemical staining for GLI1 in SHH-MB PDOX brain sections showed higher USP37 levels to be associated with higher GLI1 protein levels (Figs. 2A and 5C). CCND1, a known GLI1-target was also elevated in high USP37/high-GLI1 expressing tumors along with low levels of cleaved Caspase 3, indicating an increase in proliferation and decrease in apoptosis under these conditions (Fig 5C). Tumor sections from *Ptch^+/-^* SHH-MB mice also showed a positive association between Usp37 and Gli1 protein levels (Fig 5D).

**Figure 5.**
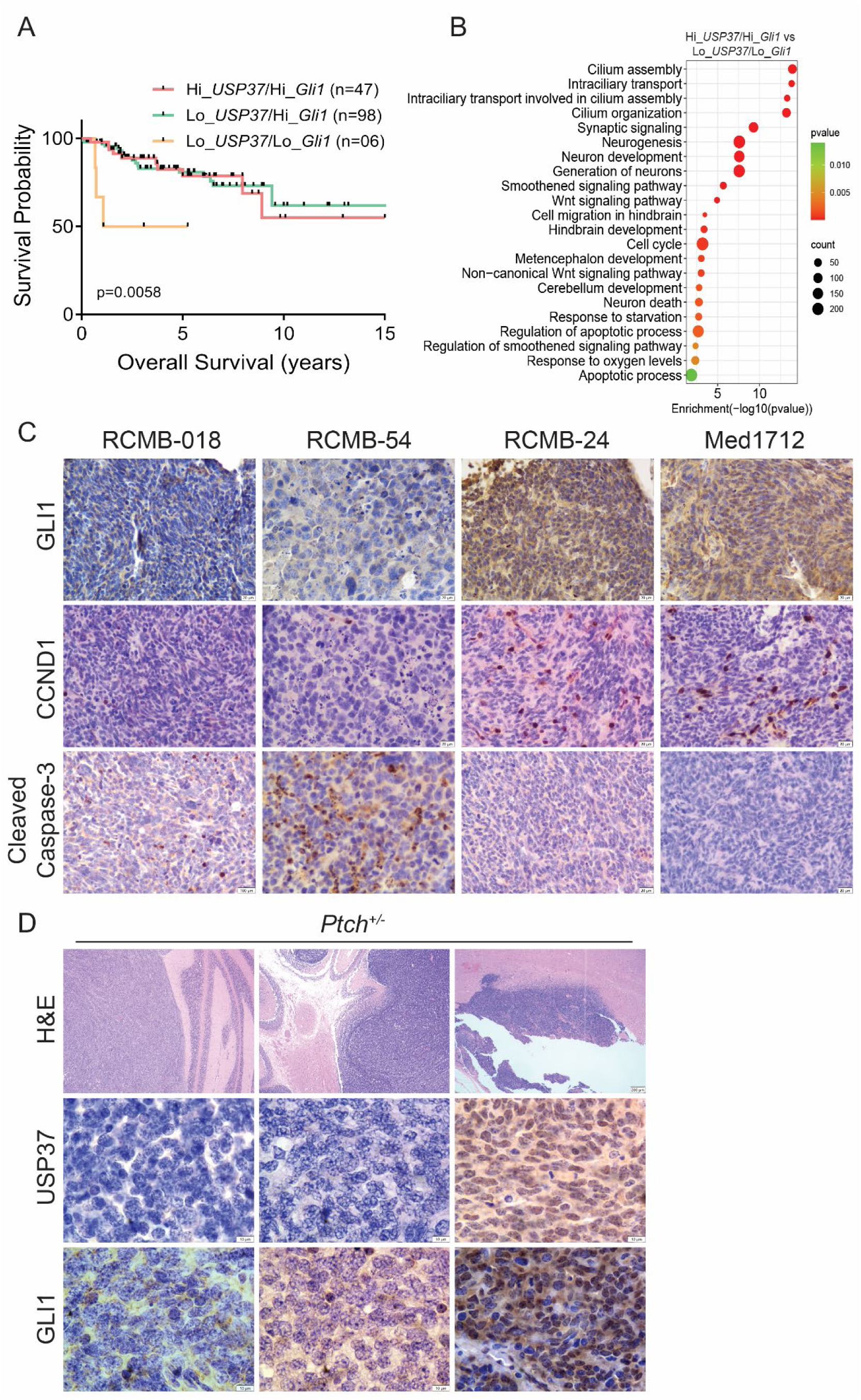
High-USP37 and high-GLI1 expression in SHH-MBs correlates with poor overall patient survival. **(A)** Kaplan–Meier plot to show a significant correlation between reduced overall survival of SHH-MB patients with high expression of *USP37* and *GLI1* in their tumors (GSE85217) [70]. (B) Pathway analysis to show significantly enriched pathways in tumors with high-*USP37*/high-*GLI1* compared to low-*USP37/*high-*GLI1* expressing samples in the GSE124814 dataset [71]. IHC staining of cerebellar sections from **(C)** mice with patient-derived orthotopic xenografts and **(D)** tumor-bearing *Ptch^+/−^/REST^TG^* animals (n=3) for GLI1, CCND1, Cleaved caspase3, Usp37 and Gli1 (scale bar: 200µm for H&E (D) and 20µm for IHC).

Next, biochemical analyses were carried out to assess if GLI1 is a target of USP37 in the neural tumor, MB. In the ONS76 and UW228 cells GLI1 levels were reduced in the presence of CHX, which could be rescued in the presence of MG132, suggesting that GLI1 is subject to proteasomal degradation, likely in a ubiquitination-mediated process (Fig 6A). Next, co-immunoprecipitation assays were done to show a strong interaction of endogenous USP37 with GLI1 in UW228 and ONS76 cells (Figs. 6B). Overexpression of USP37 in both SHH-MB cell lines promoted an increase in the levels of GLI1 and its targets, N-MYC and CCND1 (Fig 6C). *In vivo* DUB assays showed an increased GLI1 levels at 16 and 24 h post-transient transfection of WT but not the catalytically impaired *USP37* (*USP37^C350^*^S^) mutant in ONS76 (Fig. 6D). In UW228 cells, GLI1 levels were increased at 48 h post-transgene induction (Fig. S3A). However, a decrease in GLI1- associated high molecular weight bands was detected at 16 and 24 h (Fig. S3B). Anti-FLAG antibody was used to confirm expression of both *USP37* transgenes. Histone H3 served as the loading control (Fig. 6D). Lastly, *in vitro* DUB assays were performed using tagged proteins affinity purified from transiently transfected 293 cells to assess if GLI1 is a direct target of USP37. Incubation with WT USP37 caused a strong increase in the laddering pattern of ubiquitinated-GLI1 protein as well as GLI1 protein at the expected molecular weight of ∼150 KDa (Fig. 6D). Increase in low molecular weight ubiquitin (∼10 KDa) was also seen following incubation with WT USP37 (Fig. 6D). In contrast, incubation with USP37^C530S^, USP1 or WT-USP37 in the presence of NEM did not support GLI1 stabilization (Fig. 6D). The above data confirm that GLI1 protein levels in SHH-MB cells is controlled by USP37-mediated deubiquitination.

**Figure 6.**
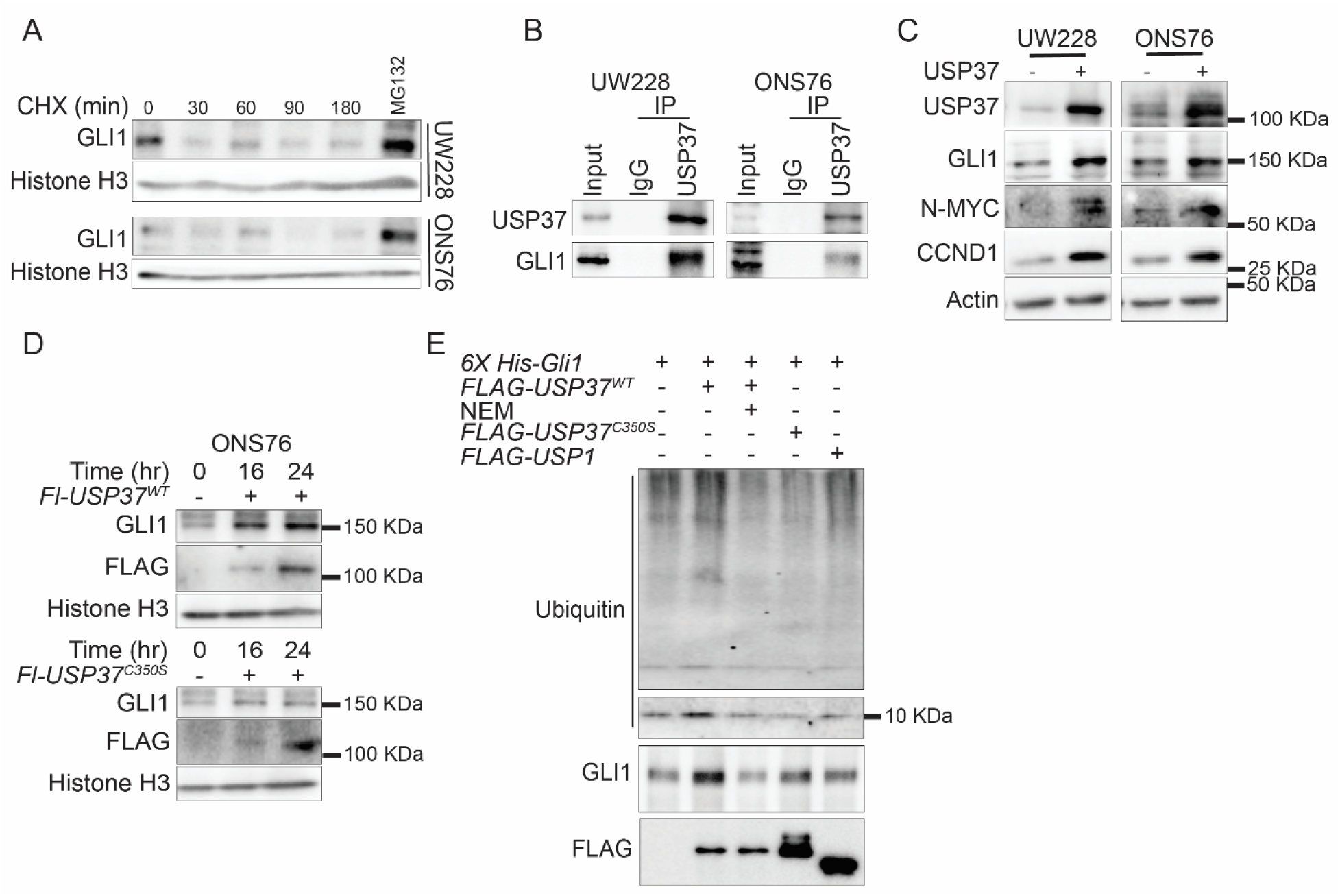
GLI1 is a target of USP37 in SHH-MB. **(A)** Western blot analyses to show changes in the levels of GLI1 in UW228 and ONS76 cells after treatment with CHX for 0 min to 180 min. MG132 was added to a second sample treated with CHX to show proteasomal control of GLI1 levels. **(B)** Co-immunoprecipitation assay using control IgG or anti-USP37 antibody to show interaction between endogenous USP37 and GLI1 proteins in UW228 and ONS76 cells. **(C)** Western blot analysis to demonstrate an increase in the levels of GLI1 and its target gene products (NMYC and CCND1) in UW228 and ONS76 cells following overexpression of transiently transfected USP37. **(D)** *In vivo* DUB assay to show a longitudinal increase in GLI1 protein levels after transient expression of *Fl-USP37^WT^* but not with *Fl- USP37^C350S^* mutant proteins in ONS76 cells. Overexpressed USP37 proteins are shown using anti- FLAG antibody. Histone H3 served as a loading control. **(E)** *In vitro* DUB assay was performed by co-incubation of purified HA-Ub-6XHis-GLI1 with Fl-WT-USP37 (lane 2), Fl-WT USP37 in the presence of NEM (lane3), Fl-USP37^C350S^ (lane 4) and Fl-USP1 (lane 5). The reaction containing purified HA-Ub-6XHis-GLI1 alone is shown in lane 1.

## Discussion

DUBs are a class of catalytic enzymes that remove ubiquitination moieties from proteins to regulate their stability and or function [80]. Deregulation of DUBs is linked to critical cellular processes such as cell cycle control, apoptosis, and DNA repair, contributing to the development and progression of various cancers [81, 82]. USP37 is a cysteine protease that has been reported to control biological processes such as chromosomal cohesion and mitotic progression, cell cycle progression, stemness and cancer cell therapeutic sensitivity, metastasis, DNA damage response and epithelial to mesenchymal transition [58, 60, 63, 65, 67, 68, 83]. Our previous work had implicated USP37 in the stabilization of the CDKi–p27 and showed that USP37 downregulation in REST-driven SHH-MBs drives cell proliferation and prevents cell cycle exit [35]. In follow up studies, we demonstrated that REST-associated G9a histone methyltransferase was involved in the epigenetically silencing of *USP37* in REST-high SHH-MB cells [36]. Overall, these studies suggested a tumor suppressive role for USP37 in REST-driven SHH-MBs. However, in non-neural cells, USP37 was also shown to exhibit oncogenic properties [61, 62, 84, 85], which suggested that USP37 may have cell and context-specific tumor suppressive and oncogenic roles.

In the current study, the identification of Raptor as a novel target of USP37 links it to the mTORC1 complex, and through it to the regulation of protein translation and potential management of cellular energetics and metabolism [86–88]. mTORC1 directly controls protein translation through its kinase activity and phosphorylation of S6 kinase (S6K) and 4EBP1, and downregulation of their activity under conditions of low *USP37* would be expected to significantly impact cap-dependent protein translation [87]. However, REST elevation in SHH-MB cells is associated with increased cell proliferation, which requires protein translation [39, 48]. This then raises questions on how these cancer cells meet the demand for new protein synthesis associated with rapidly dividing cells. A possible scenario is that in context of REST elevation and USP37 downregulation forces SHH-MB cells to switch to non-canonical cap-independent protein translation initiation mechanisms to meet the increased cellular demand for proteins necessary for stress adaptation and survival [89, 90]. Indeed, MB cells have been shown to employ internal ribosome entry site (IRES) elements to initiate protein translation without relying on the 5’ cap structure, and to selectively elevate the translation of oncoproteins essential for tumor cell growth and survival [91, 92]. However, additional studies are needed to demonstrate that indeed USP37-deficient cells rely on non-canonical translational initiation mechanisms, as well as identify the specific molecular mechanisms deployed under conditions of low mTORC1 activity.

Impaired mTORC1 complex activity is also known to reduce anabolic processes such as protein and lipid biosynthesis and decrease translation of mitochondrial proteins and biogenesis to modulate cellular energy production [87, 88]. Under these conditions, cells exhibit an increased reliance on catabolic and scavenging mechanisms such as autophagy for survival [93, 94]. In fact, in previous work we showed that REST elevation in SHH-MB cells upregulates HIF1a expression and autophagy [40]. Interestingly, in earlier work we also identified a role for REST in the phosphorylation of AKT, tumor cell infiltration and leptomeningeal dissemination [48]. AKT is also shown to modulate mTORC1 complex activity through TSC1/2 and PRAS40 [95, 96]. Thus, it is possible that *REST* elevation and consequent silencing of *USP37* expression may allow the survival of infiltrative and metastatic tumor cells through engagement of the cap-independent protein translation machinery as well as co-opting catabolic processes. These data point to a role for USP37 in the adaptation of REST-driven SHH MBs to cell stress, allowing for cell survival.

In contrast, in the absence of perturbations in REST levels in SHH-MBs or when USP37 expression is elevated and GLI1 is stabilized in a USP37-dependent manner, the upregulation of GLI1 target oncogene products such as *N-MYC* and *CCND1* drives proliferation of SHH-MB cells [97]. Work from other groups has shown that GLI1 is subject to both transcriptional and post-translational control by ubiquitination [98–101]. Our findings are also consistent with work by Qin et al, which showed USP37-mediated increase in GLI1 stability in breast cancer cells [67]. Although not tested, it may also facilitate an auto feedback loop to increase *GLI1* transcription. Here, USP37 appears to function as an oncogene. Paradoxically, when *REST* is elevated and *USP37* is downregulated, and presumably GLI1 levels are reduced, SHH signaling continues to be maintained by mechanisms that are yet to be elucidated. GLI1 is one of three members of the GLI family of transcription factors, and it is possible that redundancy between GLI1 and GLI2 supports SHH pathway activity [102, 103]. It is also important to note that USP37 is not the only GLI1-specific DUB and HAUSP/USP7, USP48 and USP21, which have been identified as GLI1 deubiquitinating enzymes in non-MB systems, and whether these compensate for *USP37* loss to allow proliferation of REST-high SHH-MBs remains to be investigated [104–106]

This work also highlights the paradoxical duality of USP37 function in medulloblastoma. This is not unique to USP37 since contradictory reports exist in the literature on the functional duality of proteins as tumor promoters or suppressors [107]. SHH-MBs tumors have their origins in cerebellar granule precursor cells (CGNPs), and potentially cells at different stages of neuronal lineage commitment [108, 109]. REST elevation in these different neuronal lineage commitment states may give rise to the clusters of SHH-MB cells described in Fig. 1D and create unique cellular contexts in which USP37 interaction with various substrate(s) may occur. Alternatively spliced isoforms of USP37 are also suggested to exist, and their preferential expression under normal or elevated REST expression conditions may be an additional avenue for context-specific USP37-substrate interactions. These studies are being actively investigated in our laboratory.

Our findings described here support a growing body of literature that suggest that non-canonical activation of GLI1 may turn on SHH pathway signaling independent of SMO activation [98, 110, 111]. Critically, tumors (including MBs), with non-canonical activation of GLI signaling are less sensitive to SMO inhibitors [112–114]. GLI inhibitors are under consideration for other cancers, yet targeting GLI proteins is thought to be challenging. GANT61, which interferes with GLI-DNA binding, is the most promising of GLI antagonists, but its clinical use is restricted by its pharmacological properties [115]. SRI-38832, a derivative of GANT61 is more potent, but is not commercially available [116]. ATO, which is FDA-approved against promyelocytic leukemia, has shown GLI inhibition in mouse models of MB and basal cell carcinoma [117]. If effective, GLI inhibitors may have promise against USP37-high SHH-MBs. While G9a is involved in the silencing of *USP37* expression, the mechanisms underlying *UPS37* upregulation in SHH-MBs are unclear and defining this process may allow for further therapeutic advances. However, the duality of USP37 function in SHH-MBs (Fig. 7), which our work clearly highlights, must be taken into consideration while effective therapeutic strategies are being developed.

**Figure 7.**
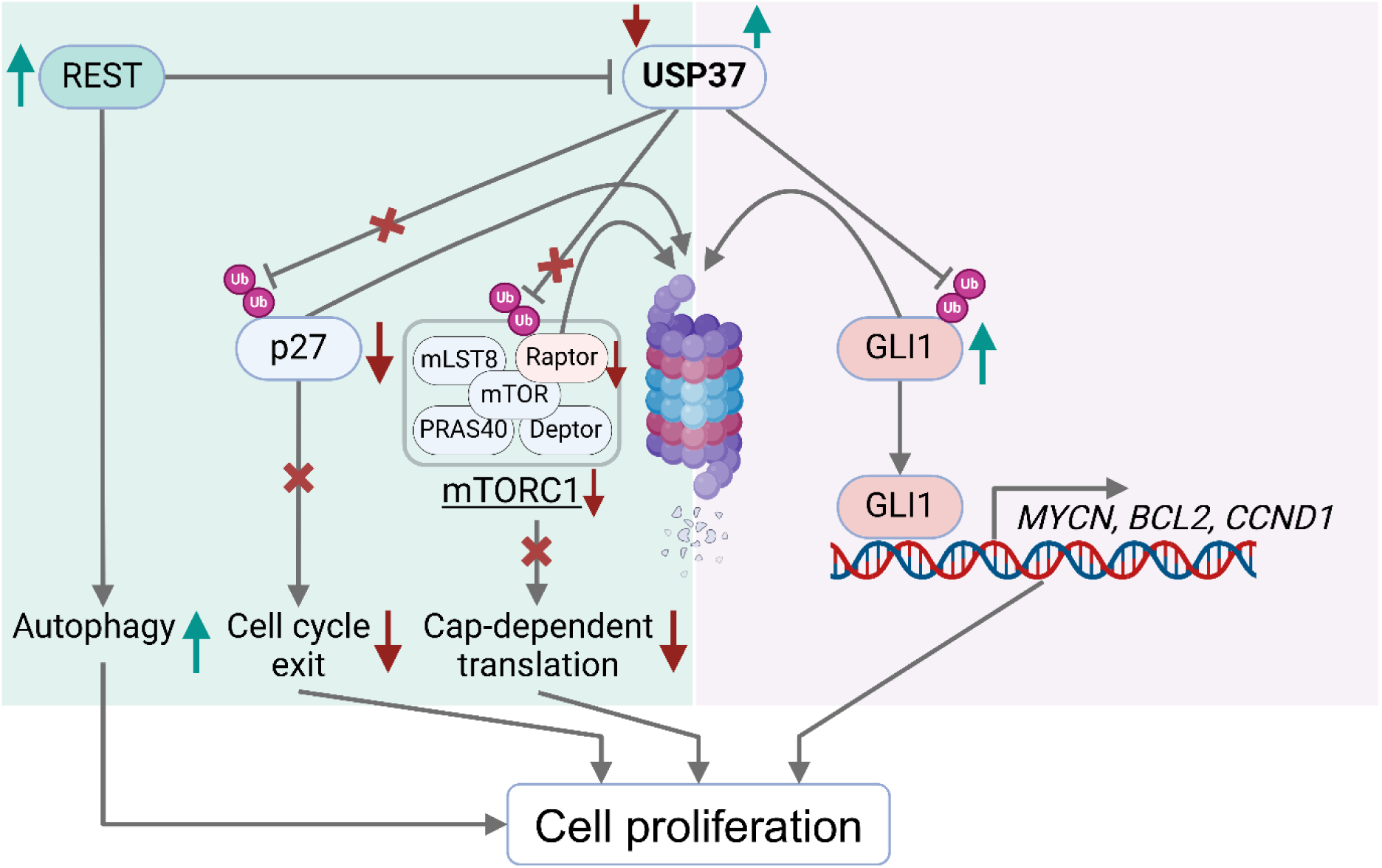
**Model to explain the context-specific role of USP37 in SHH-MB cells**. Under high REST conditions, USP37 is downregulated, leading to the destabilization of Raptor and subsequent inactivation of mTORC1. This inactivation decreases cap- dependent protein translation initiation. Conversely, when USP37 levels are elevated, it stabilizes the GLI1 protein to promote proliferation of SHH-MB cells.

## Acknowledgements

The work was supported by grants through the National Institutes of Health (NIH) (5R01NS079715) and the Addis Faith Foundation to VG.

## Author contributions

AS, DC, AH and VG contributed to design, execution, and interpretation of experiments. YY, TD and VR also performed experiments. AS, AH, TD and VG wrote the manuscript. VG provided funding support.

## Data availability statement

All transcriptomic analyses were performed using previously published datasets: GSE85217 (https://www.ncbi.nlm.nih.gov/geo/query/acc.cgi?acc=GSE85217) and GSE124814 (https://www.ncbi.nlm.nih.gov/geo/query/acc.cgi?acc=GSE124814).

## Supplementary Information

**Figure S1.**
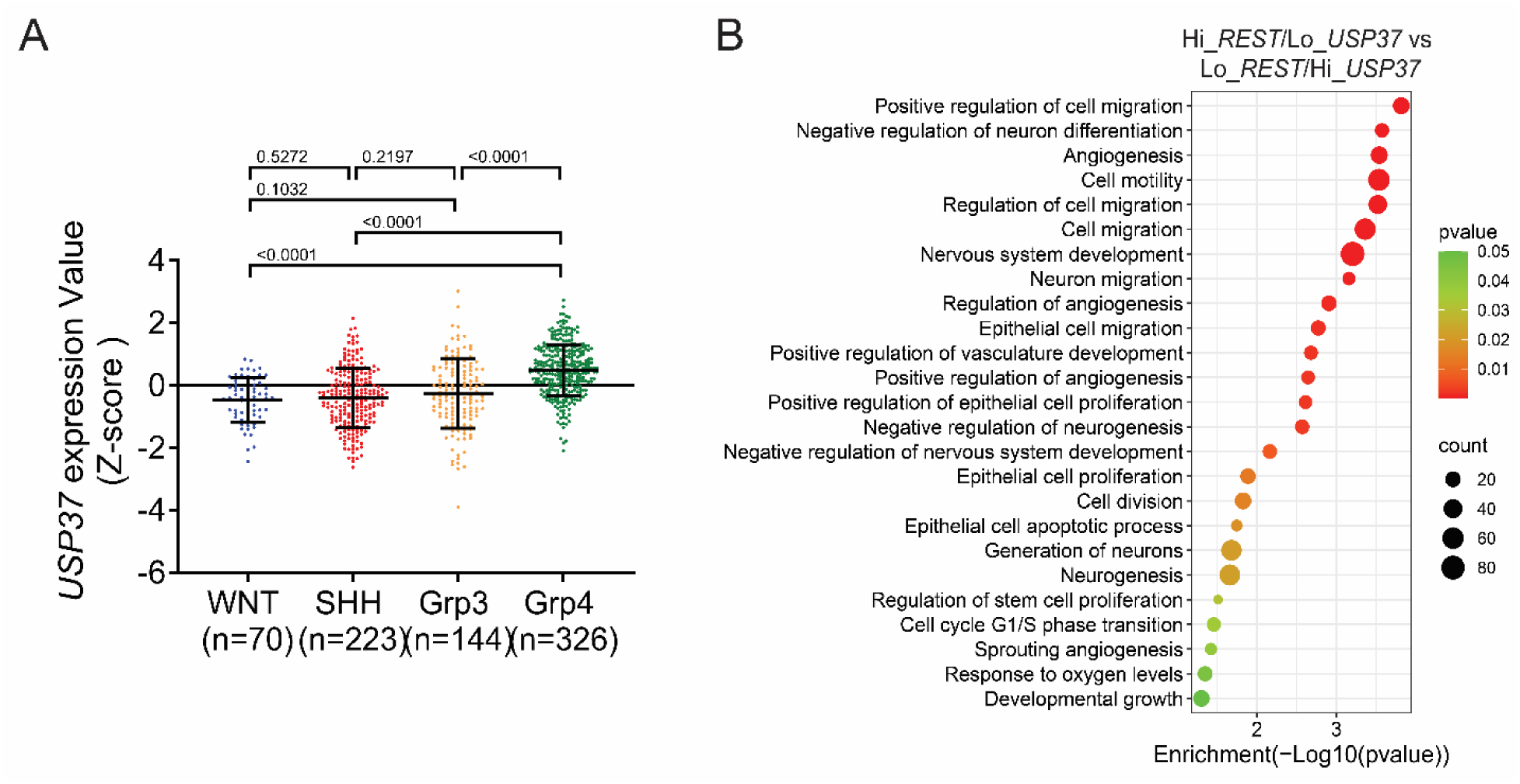
**(A)** Dot plot showing the expression of *USP37* in the four MB subgroups. Microarray dataset GSE85217 [66] was used for the analysis. **(B)** Significantly enriched pathways in tumors with high-*REST*/low-*USP37* expression relative to samples with low- *REST*/high-*USP37* expression in GSE85217 dataset [66].

**Figure S2.**
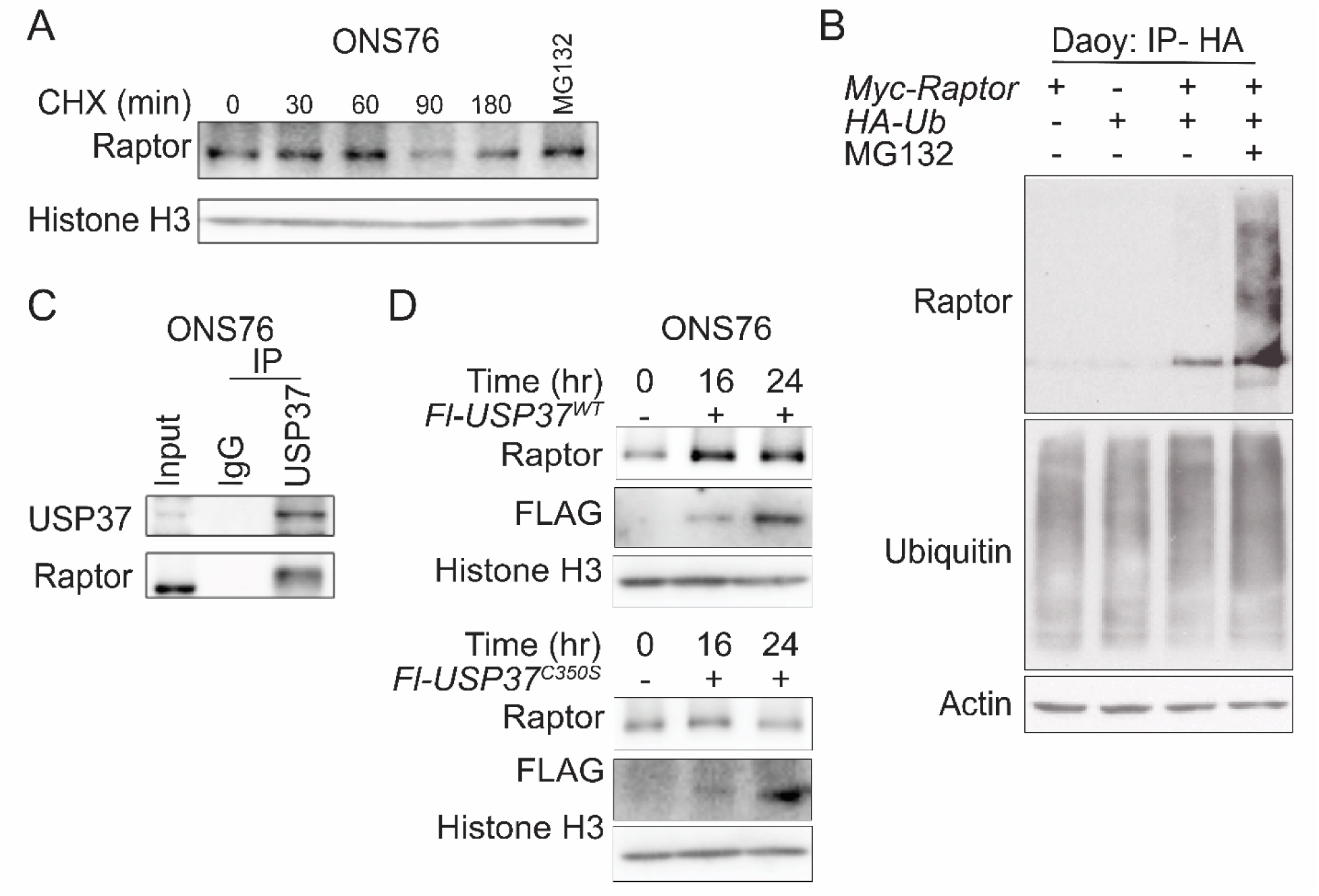
**(A)** Western blot showing post-transcriptional and proteasomal control of Raptor in ONS76 cells treated with Cycloheximide (CHX) (lanes 1-5) alone and with MG132 (lane 6). Co-immunoprecipitation assay showing **(B)** Raptor ubiquitination in DAOY cells transiently transfected with Myc-tagged Raptor and HA-tagged ubiquitin. **(C)** Interaction between endogenous USP37 and Raptor in ONS76 cells. Anti-USP37 antibody or control IgG were used. **(D)** *In vivo* DUB assay using transiently transfected FLAG (Fl)-tagged wildtype (WT) USP37 (top panel) and mutant USP37 (USP37^C350S^) (bottom panel) in ONS76 cells to assess changes in the levels of endogenous Raptor.

**Figure S3.**
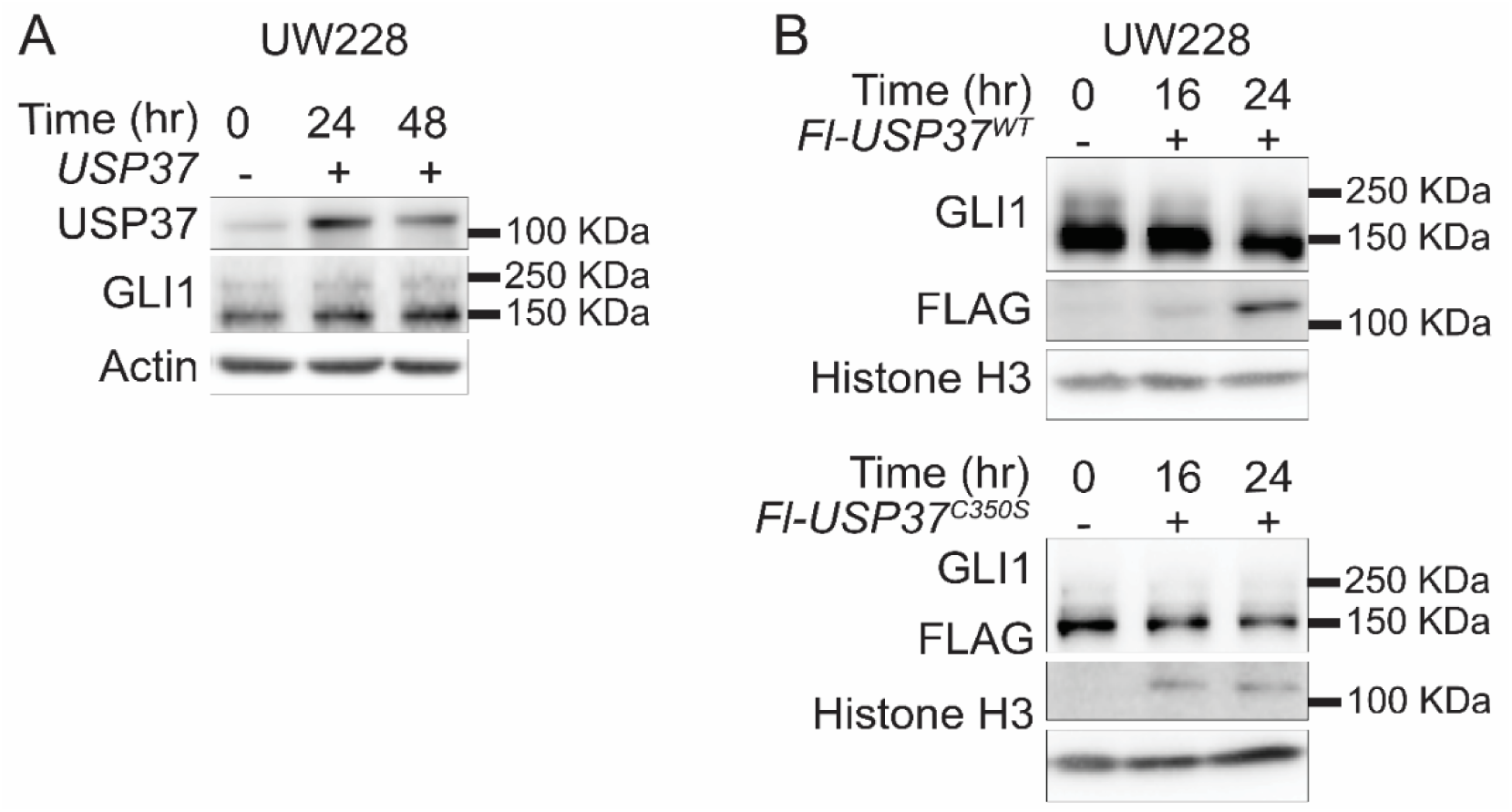
**(A)** Western blot showing the level of GLI1 in UW228 cells with USP37 overexpression over 48 h. **(B)** *In vivo* DUB assay to show decrease in GLI1 protein after transient expression of *Fl-USP37^WT^* but not with *Fl- USP37^C350S^* mutant proteins in UW228 cells. Overexpressed USP37 proteins are shown using anti- FLAG antibody. Histone H3 served as a loading control.

**Table S1:** List of antibodies used in this study.

| **Antibody** | **Company** | **Catalogue number** |
| --- | --- | --- |
| **Western blot** |  |  |
| Akt | Cell Signaling Technology | 9272 |
| Phospho-Akt (Ser473) | Cell Signaling Technology | 3787 |
| GSK-3β | Cell Signaling Technology | 9315 |
| Phospho-GSK-3β (Ser9) | Cell Signaling Technology | 9323 |
| p27 | BD Biosciences | 610242 |
| Phospho-p27 (T157) | Abcam | ab85047 |
| Rictor | Novus Biologicals | NB100-612 |
| Phospho-Rictor (Thr1135) | Cell Signaling Technology | 3806 |
| mTOR | Cell Signaling Technology | 2983 |
| MLST8 (GβL) | Cell Signaling Technology | 3274 |
| mSIN1 | ThermoFisher Scientific | A300-910A |
| Raptor | Cell Signaling Technology | 2280 |
| Phospho-Raptor (Ser792) | Cell Signaling Technology | 2083 |
| PRAS40 | Cell Signaling Technology | 2691 |
| Deptor | Novus Biologicals | NBP1-49674 |
| 4EBP1 | Cell Signaling Technology | 9644 |
| Phospho-4E-BP1 (Thr37/46) | Cell Signaling Technology | 2855 |
| USP37 | Abcam | ab190184 |
| eIF4G | Cell Signaling Technology | 2498 |
| eIF4E | Cell Signaling Technology | 2067 |
| Gli1 | Abcam | ab134906 |
| N-Myc | Abcam | ab16898 |
| CCND1 | Cell Signaling Technology | 55506 |
| Ubiquitin | Cell Signaling Technology | 3933 |
| Flag | Millipore Sigma | F1804 |
| Histone H3 | ThermoFisher Scientific | PA5-16183 |
| Actin | Millipore Sigma | A1978 |
| **Immunohistochemistry** |  |  |
| Ki67 | Abcam | ab15580 |
| TUBB3 | Abcam | ab18207 |
| REST | Abcam | ab202962 |
| USP37 | ThermoFisher Scientific | PA5-100697 |
| CCND1 | Cell Signaling Technology | 55506 |
| Raptor | Cell Signaling Technology | 2280 |
| Gli1 | ThermoFisher Scientific | MA5-32553 |
| CCND1 | Cell Signaling Technology | 55506 |
| Cleaved caspase 3 | Cell Signaling Technology | 9664 |
| **Co-immunoprecipitation** |  |  |
| USP37 | Proteintech | 18465-1-AP |
| 4EBP1 | Cell Signaling Technology | 9644 |
| eIF4E | Cell Signaling Technology | 2067 |

